# Steady-state responses in the somatosensory system interact with endogenous beta activity

**DOI:** 10.1101/690495

**Authors:** Michel J. Wälti, Marc Bächinger, Kathy L. Ruddy, Nicole Wenderoth

## Abstract

Brain oscillations have been related to many aspects of human behavior. To understand a potential causal relationship, it is of great importance to develop methods for modulating ongoing neural activity. It has been shown that external rhythmic stimulation leads to an oscillatory brain response that follows the temporal structure of the presented stimulus and is assumed to reflect the synchronization of ongoing neural oscillations with the stimulation rhythm. This interaction between individual brain activity and so called steady-state evoked potentials (SSEPs) is the fundamental requirement of neural entrainment. Here, we investigate whether neural responses to rhythmic vibrotactile stimulation, measured with EEG, are dependent on ongoing individual brain oscillations, and therefore reflect entrained oscillatory activity. For this, we measured phase synchronization in response to rhythmic stimulation across various frequencies in the alpha and beta band. Three different stimulation intensities were applied for each frequency relative to the individual sensory threshold. We found that a higher stimulation intensity, compared to lower intensities, resulted in a more pronounced phase synchronization with the stimulation signal. Moreover, EEG responses to low stimulation frequencies closer to individual beta peak frequencies revealed a higher degree of entrainment, compared to stimulation conditions with frequencies that were more distant to endogenous oscillations. Our findings provide evidence that the efficacy of vibrotactile rhythmic beta stimulation to evoke a SSEPs is dependent on ongoing brain oscillations.

## Introduction

During cognitive tasks brain rhythms of different frequencies can be recorded at the scalp using electroencephalography (EEG), that are correlated with cognitive and behavioral processes. In recent years, it has been suggested that various non-invasive methods, such as repetitive transcranial magnetic stimulation (rTMS) or transcranial alternating current stimulation (tACS) are capable of frequency-specific modulation of brain rhythms. Hence, they have been used to test the causal relationship between the strength of neuronal oscillations and behavior (for a review, see Thut, Schyns, & Gross, 2011). However, the interaction of these externally applied magnetic or electric fields with ongoing brain rhythms is still poorly understood. An alternative form of frequency-specific modulation of brain rhythms can be achieved with rhythmic sensory stimulation which induces steady-state evoked potentials (SSEPs) following the temporal frequency of the driving stimulus (Regan, 1977). SSEPs are usually measured with electroencephalography (EEG) and have been documented in the visual (steady-state visually evoked potentials, SSVEPs), the auditory (auditory steady-state responses, ASSRs) and the somatosensory (steady-state somatosensory evoked potentials, SSSEPs) systems (for a review, see Vialatte, Maurice, Dauwels, & Cichocki, 2010). However, the underlying mechanism of SSEPs, is still debated (see Zoefel, Ten Oever, & Sack, 2018). While some studies report evidence for entrainment of ongoing brain oscillations by showing an interaction between stimulation and endogenous activity (Notbohm, Kurths, & Herrmann, 2016; Schwab et al., 2006), others found responses to be nothing more than regular repetitions of evoked neural potentials (Capilla, Pazo-Alvarez, Darriba, Campo, & Gross, 2011; C. Keitel, Quigley, & Ruhnau, 2014). In contrast to simply overlaying evoked potentials, entrainment modulates brain oscillations. To test whether brain responses to rhythmic sensory stimulation reflect entrainment, it is therefore necessary to provide evidence of a dependency on ongoing brain oscillations. This dependency arises from theoretical considerations and has been the main research interest in studies supporting the entrainment hypothesis (see Zoefel et al., 2018). The so-called Arnold tongue describes a classical model of entrainment between two oscillators depending on stimulation rhythm and intensity (Pikovsky, Rosenblum, Zaks, & Kurths, 1999). The model predicts that a high stimulation intensity evokes a larger response than a low intensity but also entrainment to be most pronounced when the stimulation frequency matches the frequency of the endogenous system (e.g. Fröhlich, 2015). In a recent study, Notbohm et al. (2016) reported that the coupling of EEG oscillations with rhythmic visual flicker follows the characteristics of the Arnold tongue. They showed that the EEG response to the stimulation depends on both the stimulation intensity and the distance between the flicker frequency and the participants’ individual alpha frequency (IAF; Notbohm et al., 2016). Other studies found similar effects for tACS over occipital cortex which either increases EEG power at participants’ visual alpha frequency (Helfrich et al., 2014; Zaehle, Rach, & Herrmann, 2010), or biases ongoing alpha oscillations in visual cortex towards the exogenous stimulation frequency (Minami & Amano, 2017). Most previous studies have tested entrainment in visual cortex targeting the alpha rhythm. Here we test whether it is also possible to entrain endogenous brain rhythms of primary sensorimotor cortex via rhythmic tactile stimuli. It is well known that the sensorimotor cortex exhibits strong activity in the alpha (8 – 12 Hz) and beta band (15 – 30 Hz) when subjects are at rest (Crone et al., 1998; Fransen, Dimitriadis, van Ede, & Maris, 2016). During sensorimotor processing and, particularly, during motor execution, both rhythms desynchronize (Bauer, Oostenveld, Peeters, & Fries, 2006; Kilavik, Zaepffel, Brovelli, MacKay, & Riehle, 2013; Spitzer & Haegens, 2017) but resynchronize quickly once the movement is completed, implicating both rhythms in sensorimotor processing (Kilavik et al., 2013; Pfurtscheller & Lopes da Silva, 1999; Ploner, Gross, Timmermann, Pollok, & Schnitzler, 2006; Rajagovindan & Ding, 2011). Here we ask whether the alpha and the beta rhythms in primary sensorimotor cortex are equally susceptible to entrainment via tactile stimuli. We hypothesized that repetitive tactile stimulation at or close to the intrinsic alpha and beta frequency peaks of the sensorimotor system should be more effective than stimulation at other frequencies. To that end, we probed whether characteristics of the Arnold tongue can be found when measuring phase synchronization in response to vibrotactile stimulation at different frequencies in the alpha and beta bands. Three different stimulation intensities were applied for each frequency which were defined relative to individual sensory threshold. We expected steady-state EEG responses to be more pronounced for (i) high-intensity stimulations compared to low intensities, and (ii) at stimulation frequencies closer to individual alpha (IAF) and/or beta frequencies (IBF) compared to frequencies more distant from endogenous oscillations.

## Materials and methods

### Participants

25 participants took part in the experiment in exchange for monetary compensation (20 Swiss francs per hour). The participants had no reports of psychiatric disorders and were all right-handed according to the Edinburgh Handedness Inventory (Oldfield, 1971). The study protocol was approved by the local ethics committee and was conducted in accordance with the Declaration of Helsinki. All participants provided written consent. 3 participants were excluded from the analysis due to technical problems during EEG recording or because they did not follow the instructions (e.g. falling asleep during the EEG recording), resulting in a final sample of 22 participants (female: 8; age: range = 18 – 29, M ± SD = 22.8 ± 2.6).

### Design and procedure

The experiment was conducted in two parts. In a first part, we determined participants’ IAF and IBF in a simple finger tapping task. The second part consisted of the main experiment with different conditions of vibrotactile stimulation to participants’ right index finger. Throughout both parts, EEG was recorded (see below).

#### Part I: Detection of IAF and IBF

Participants were comfortably seated in a dark sound-attenuated room, approximately 100 cm away from a computer screen (27-inch) and were instructed to execute a self-paced finger tapping task with their right hand, in which they tapped and gently squeezed their thumb against each of the other fingers successively. One run consisted of an alternating green and red fixation cross appearing in the center of the screen (each 9 times for 10 seconds). Green indicated that the participant should execute the finger tapping, and red indicated that they should rest their hand. Each participant performed two runs.

#### Part II: Vibrotactile stimulation

During the main experiment, participants’ right index finger was stimulated with a rhythmic vibrotactile stimulation. For the stimulation, an earphone (YVE-01B-03, Yeil Electronics Co., South Korea) with a diameter of 17 mm was taped to the participants finger. The earphone was modified to produce vibrations via a coil actuator with a mass attached which vibrates when current is passed through. The stimulation signals were generated in MATLAB (The MathWorks Inc., USA) and sent to the stimulation devices via a National Instruments Card (NI-USB 6343) and an amplifier (Dancer Design, St. Helens, UK). The stimulation was elicited by driving the stimulator with a carrier frequency of 200 Hz which was frequency-modulated by a square wave corresponding to the stimulation frequency (Breitwieser, Kaiser, Neuper, & Müller-Putz, 2012). This time course of the driving voltage had a resolution of 0.1 ms. For each participant, the individual stimulation threshold was determined prior to the main experiment by applying a 20 Hz stimulation stimulus which started at sub-threshold intensity and increased in steps of 10% increments of output voltage until it was perceived. This procedure was repeated three times. The mean values of the detection thresholds indicated the individual stimulation threshold for each participant. Based on the individual stimulation threshold three stimulation intensities were set: (i) sub-threshold intensity corresponding to 0.5 x threshold intensity (sub-threshold), (ii) a low supra-threshold intensity corresponding to 20 x threshold intensity (low), and (iii) a high supra-threshold intensity corresponding to 200 x threshold intensity (high). During the main experiment (Figure 1), all stimulation frequencies (i.e. 6 – 24 Hz) were applied with the same sub-threshold, low and high intensities, respectively. Note that the range of our experimental frequencies was rather narrow such that all activate the same tactile mechanoreceptors (Meissner’s corpuscles, 5 – 50 Hz; see Johansson & Flanagan, 2009). After threshold detection, the main experiment began which consisted of 3 blocks each consisting of 10 stimulation conditions which were carried out in random order (resulting in 30 stimulation conditions in total). These 30 conditions were composed of 10 different stimulation frequencies ranging from 6 to 24 Hz in steps of 2 Hz, each administered with the three stimulation intensities (sub-threshold, low or high). Each stimulation lasted for 40 seconds and was preceded by a 10 second period without stimulation. Participants were instructed to sit as still as possible and direct their gaze at a white fixation cross centered on a black screen.

**Figure 1.**
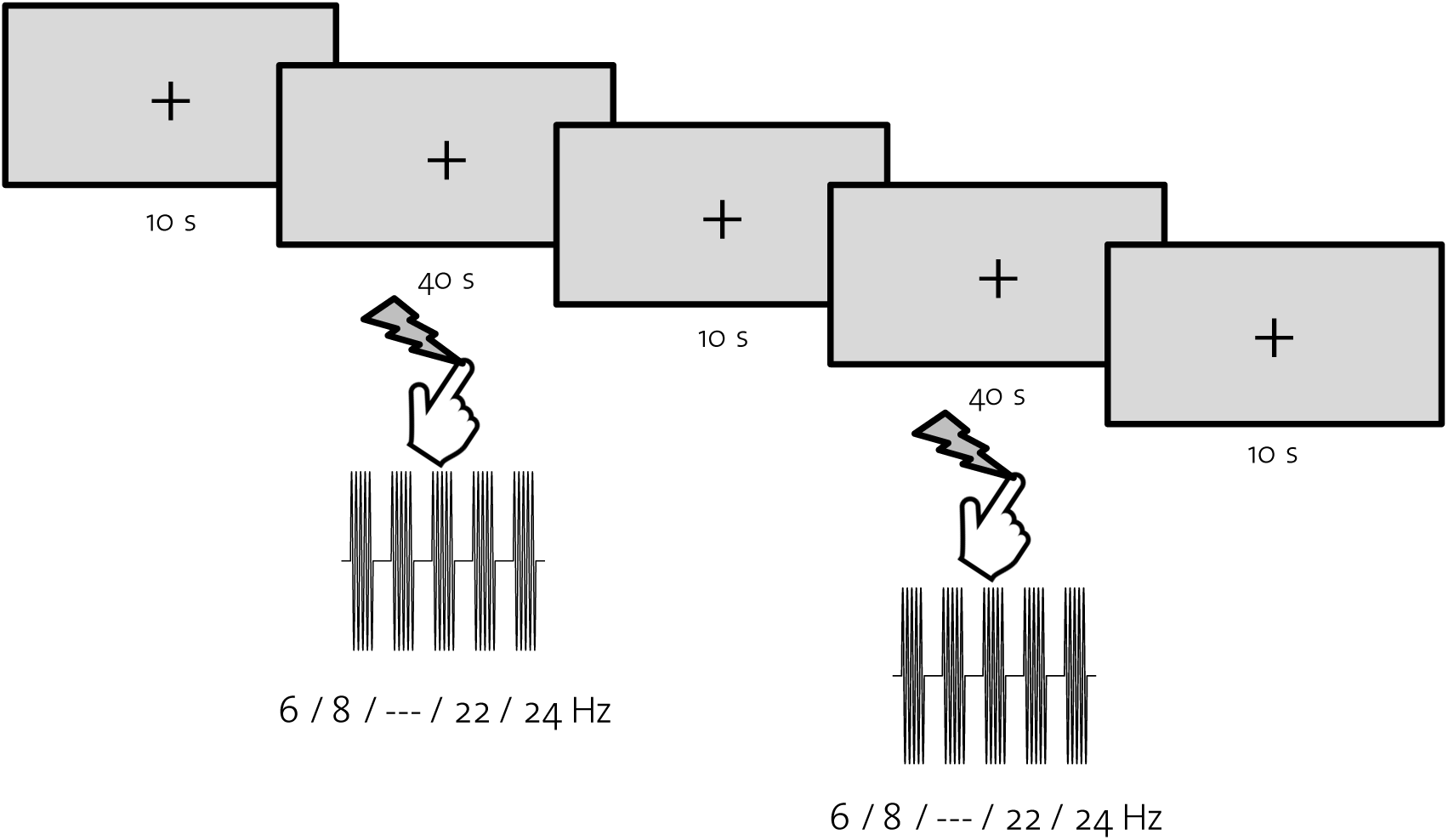
Experimental procedure. Participants’ right index finger was stimulated with frequencies ranging from 6 to 24 Hz at 3 different stimulation intensities. Each stimulation lasted for 40 s, preceded by a 10 s period without stimulation (rest).

### Data acquisition

EEG data were acquired at 1000 Hz using a 64-channel Hydrocel Geodesic EEG System (Electrical Geodesic Inc., USA), referenced to Cz (vertex), with an online Notch filter (50 Hz) and high-pass filter at 0.3 Hz. Impedances were kept below 50 kΩ.

### Preprocessing of EEG data

EEG data for both parts of the experiment were preprocessed and analyzed offline. Data were cleaned (detection and interpolation of bad electrodes), band-pass filtered (5 – 25 Hz) and further processed using independent component analysis (ICA). Artifact components (ICs) were automatically detected and removed from the data using a custom built toolbox (see Liu, Ganzetti, Wenderoth, & Mantini, 2018).

### EEG data analysis

#### Part I: IAF and IBF

To detect IAF and IBF, preprocessed data were epoched regarding the two conditions (movement and rest; each from +1 s to +9 s after start of the condition) and analyzed using fast Fourier transform (FFT). For each subject, frequency-band specific power differences in the alpha (8 – 12 Hz) and beta band (15 – 25 Hz) were calculated between movement and rest to detect event-related de-synchronization (ERD) that typically appears during movement as well as event-related synchronization (ERS) immediately after movement. Power changes were determined as percentage change from rest versus movement (Pfurtscheller & Lopes da Silva, 1999):

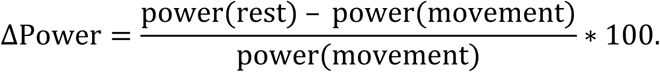

The electrode(s) exhibiting the strongest ΔPower over the contralateral hemisphere (right hand, left hemisphere) for the alpha and beta band, respectively, was selected and we determined (i) IAF as the frequency with the biggest power increase between 6 and 14 Hz (range of the stimulation frequencies); and (ii) IBF as the frequency with the most pronounced power increase between 16 and 24 Hz. Similar to before, we defined power increases as percentage change in power in the rest condition compared to the move condition (ΔPower) and normalized the resulting power spectrum to the power at the alpha or beta peak respectively.

#### Part II: Steady-state evoked potentials

In order to evaluate general effects of vibrotactile stimulation on EEG signals, we measured frequency-specific power and phase-based consistency of EEG time-series during stimulation. For this, preprocessed EEG data from the main experiment were epoched to each stimulation condition (+1 s to +39 s relative to stimulation onset). Further, time-series data were cropped in 1 s segments without overlap which were later treated as separate trials. In order to analyze steady-state responses to the stimulation frequencies we pre-defined a region of interest (ROI) consisting of nine electrodes over the somatosensory area. For each participant, we evaluated the electrode that showed the strongest SSEPs across alpha (6 Hz, 8 Hz, 10 Hz, 12 Hz, 14 Hz) and beta (16 Hz, 18 Hz, 20 Hz, 22 Hz, 24 Hz) stimulation conditions separately. For this, data averaged across trials of each high-intensity stimulation condition were transformed into its frequency components using a fast Fourier transform (FFT) and power at the corresponding stimulation frequency as well as neighboring frequencies (−2 Hz, −1Hz, +1Hz, +2Hz) were determined for each electrode within the ROI. We then calculated percentage increase of power at the stimulation frequency compared to power at neighboring frequencies for each electrode. To evaluate the electrode showing the strongest response across the alpha stimulation conditions, we averaged the calculated power changes across the corresponding conditions. The same procedure was carried out for the beta stimulation conditions. To determine stimulation effects on frequency-specific power, FFT was performed on data that had been preprocessed and averaged across trials, derived from these electrodes (strongest alpha and beta for each participant). To determine frequency-specific phase consistency of the EEG signals, we further calculated intertrial phase clustering (ITPC), which measures the extent of phase angle distributions at each time-frequency point across trials (Cohen, 2014).

#### Part II: Phase-based synchronization between EEG and stimulation signal (entrainment)

To determine entrainment, we used data from the electrodes which were earlier defined as detecting the strongest alpha and beta signals. We then measured changes in intersite phase clustering (ISPC) between the preprocessed and averaged time-series data of each stimulation condition and the corresponding stimulation signal. ISPC is a measure of phase-based connectivity between two time-series signals (Cohen, 2014). Similar to ITPC, ISPC measures phase angle distributions, with the key distinction that phase angle *differences* between two signals are measured. Similarly, to the EEG data, the stimulation signals were band-pass filtered in the range from 5 to 25 Hz and down-sampled to 1000 Hz. The single-channel EEG time-series were further band-pass filtered in the range of ±2 Hz regarding the corresponding stimulation frequency. To detect changes in phase-locked activity in response to the stimulation, we compared ISPC between the stimulation signal and the EEG signal during stimulation with ISPC prior to each condition (i.e. preceding rest period). Similar to the previously described procedure, EEG data prior to stimulation onset (−9 s to −1 s) was cropped in 1 s segments and then averaged across trials. ΔISPC was calculated by subtracting the mean pre-stimulation ISPC from the mean ISPC during stimulation for each of the 30 conditions. All scripts to execute the experiment and to preprocess and analyze EEG data were written in MATLAB (The MathWorks Inc., USA), and used functions from EEGLAB (Delorme & Makeig, 2004). Scripts using FFT, ITPC and ISPC analysis were adapted from M. X. Cohen (Cohen, 2014). For the experiment presentation code, PsychToolbox was used (Brainard, 1997).

### Statistical analysis

All statistical analyzes were performed separately for conditions targeting alpha oscillations (stimulation frequencies: 6 Hz, 8 Hz, 10 Hz, 12 Hz, 14 Hz) and beta oscillations (stimulation frequencies: 16 Hz, 18 Hz, 20 Hz, 22 Hz, 24 Hz). General effects of the stimulation signals on frequency-specific power and ITPC were analyzed using repeated-measures ANOVA models in SPSS Version 25 (IBM, USA). To test the hypothesized effects regarding phase-based connectivity between stimulation and EEG signals, we used R (R Core Team, 2018) and the lme4 package (Bates, Mächler, Bolker, & Walker, 2015) to perform linear mixed effects analyzes. We used linear mixed effects models because they account for variance arising from individual differences (Baayen, Davidson, & Bates, 2008). To ensure that observed effects are not driven by individual variability, subjects were included in the models as random factors. Fixed effects represent the influence of experimental factors on the dependent variable (here: ΔISPC). In our models, the fixed effects include stimulation intensity (sub-threshold intensity as baseline) and distance of the stimulation frequency to individual endogenous frequencies in the alpha (ΔIAF), or the beta (ΔIBF) band. Validation of our models was done by plotting residuals against fitted values.

## Results

The current study was designed to reveal variations in the effect of vibrotactile stimulation on the individual neuronal response in sensorimotor cortex. Participants were rhythmically stimulated at their right index finger (dominant hand) using three stimulation intensities and ten stimulation frequencies across the alpha and beta bands. Each stimulation lasted for 40 seconds and was preceded by a 10 second period without any stimulation.

### Detection of IAF and IBF

To detect IAF and IBF, frequency-band specific power changes due to event-related de-/synchronization (ERD/ERS) in a simple finger tapping task were determined for each subject in the alpha and beta bands. Figure 2 A/B show topographical plots and power spectra from an example subject, revealing strong power increases in the left primary motor areas (rest vs. movement) and clear peaks in the power spectra in both alpha and beta bands. Across subjects, average IAF was 11.0 Hz (± 2.1) and average IBF was 20.3 Hz (± 1.9) (see Figures 2 C/D). To evaluate whether the two spectral peaks in alpha and beta present two independent rhythms, we calculated linear correlation with the Pearson correlation coefficient between IAFs and IBFs across subjects. We found only a weak negative correlation (r = −0.22, p = 0.319) and conclude that the power changes related to the finger tapping task found in alpha and beta represent two distinct endogenous brain rhythms rather than harmonics.

**Figure 2.**
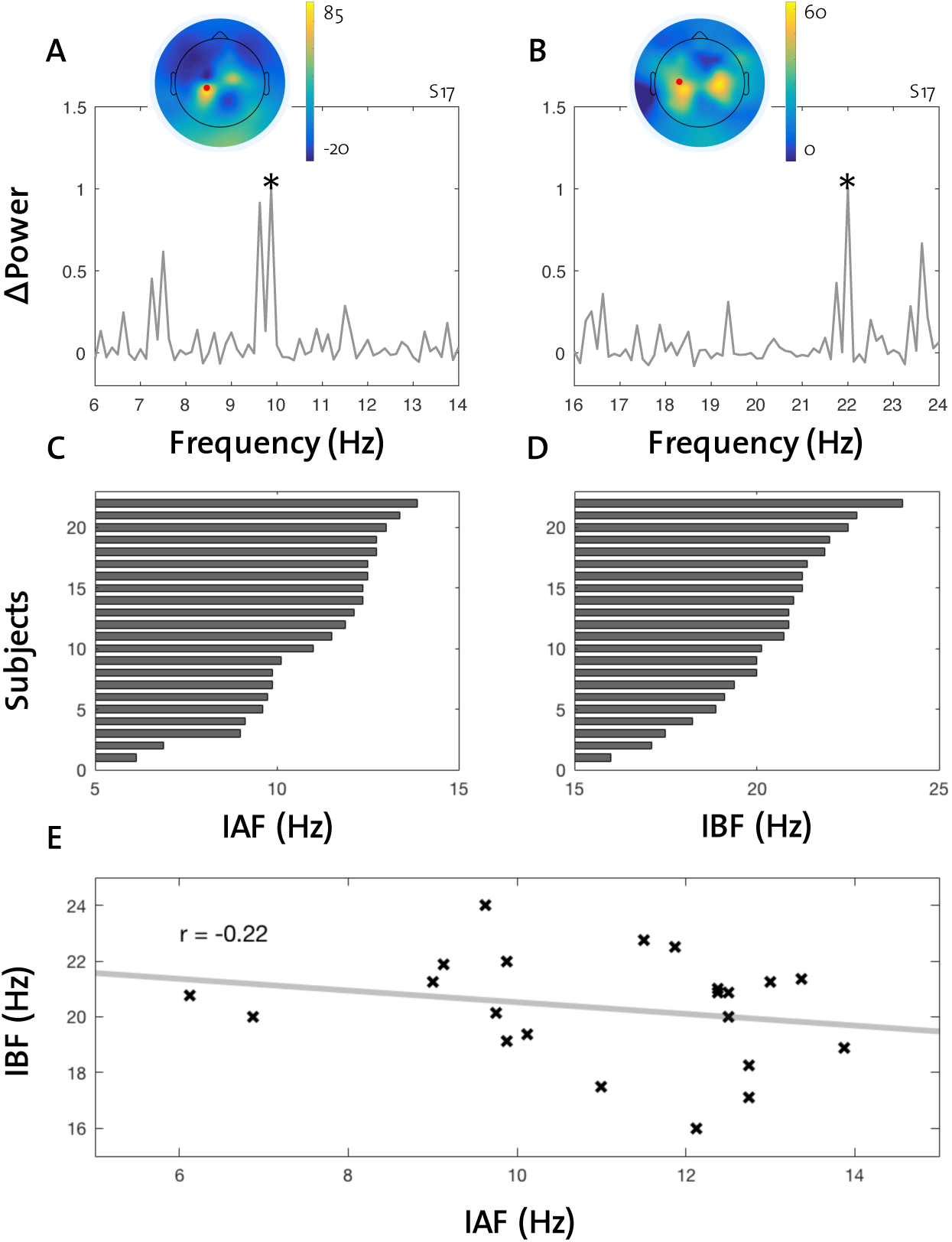
Participants’ IAF and IBF. Detection of participants’ endogenous peak frequencies in the alpha (IAF) and beta (IBF) bands by calculating frequency-band specific power changes due to event-related de-/synchronization (ERD/ERS) in a simple finger tapping task. A: Example subject’s topographical plot of ΔPower in the alpha band and power spectrum of the electrode showing the highest value (red dot). IAF was determined as frequency with the highest ΔPower (here: 9.875 Hz, marked with an asterisk). B: Same example subject’s topographical plot and power spectrum as in A, but for the beta band (IBF: 22 Hz, marked with an asterisk). C: Ordered IAF frequencies across all subjects. D: Ordered IBF frequencies across all subjects. E: Depiction of correlation with regression line between IAF and IBF values. Pearson correlation coefficient reveals only a weak negative linear dependency (r = −0.22).

### Stimulation intensity differences on steady-state evoked potentials in alpha and beta

In order to evaluate entrainment effects, we first need to determine whether our stimulation protocol reveals typical characteristics of steady-state evoked potentials over the somatosensory area across alpha and beta frequencies. Further, we are interested in general differences between the two frequency bands regarding effects of stimulation intensity. Therefore, a two-way repeated measures ANOVA with factors frequency band (alpha and beta) and stimulation intensity (sub-threshold, low, high) was used to detect general effects of different stimulation conditions on frequency-specific power and phase-clustering (ITPC) values. Figures 3 and 4 depict stimulation effects averaged across all subjects. Figure 3 shows changes in the power spectra across all stimulation conditions and Figure 4 reveals effects on ITPC. Our first analysis used power as dependent variable (derived from FFT). The results from the ANOVA reveal a main effect of stimulation intensity (F(2,20) = 50.7, p < 0.001), but not of frequency band (F(1,21) = 0.7, p < 0.411). Bonferroni adjusted post-hoc comparisons between intensities revealed a significant difference between all three levels (sub-threshold vs. low: p = 0.030; sub-threshold vs. high: p < 0.001; low vs. high: p < 0.001). Further, the interaction between frequency band and stimulation intensity show a significant effect, meaning that vibrotactile stimulations in alpha and beta have different effects on frequency-specific power across the different intensities (F(2,20) = 3.7, p = 0.041). This steeper increase in power across the stimulation intensities at beta compared to alpha (see Figure 5 A) is in line with previous research suggesting that the somatosensory system has a temporal resonance frequency in the beta band (Müller, Neuper, & Pfurtscheller, 2001; Snyder, 1992; Tobimatsu, Zhang, & Kato, 1999), thus, responding with greater amplitudes to stimulation frequencies in beta. The same approach was used to detect effects on ITPC across the two frequency bands and three stimulation intensities. Again, we found a main effect of stimulation intensity (F(2,20) = 24.8, p < 0.001), but not of frequency band (F(1,21) = 1.0, p = 0.341). Post-hoc comparisons between intensities revealed a significant difference between high intensity compared to the other two, but not between sub-threshold and low levels (sub-threshold vs. low: p = 0.293; sub-threshold vs. high: p < 0.001; low vs. high: p < 0.001). Further, no significant interaction between frequency band and stimulation intensity on ITPC was observed (F(2,20) = 2.7, p = 0.093). We conclude that rhythmic vibrotactile stimulation has similar effects on alpha and beta band frequencies regarding phase-angle distributions across trials (ITPC).

**Figure 3.**
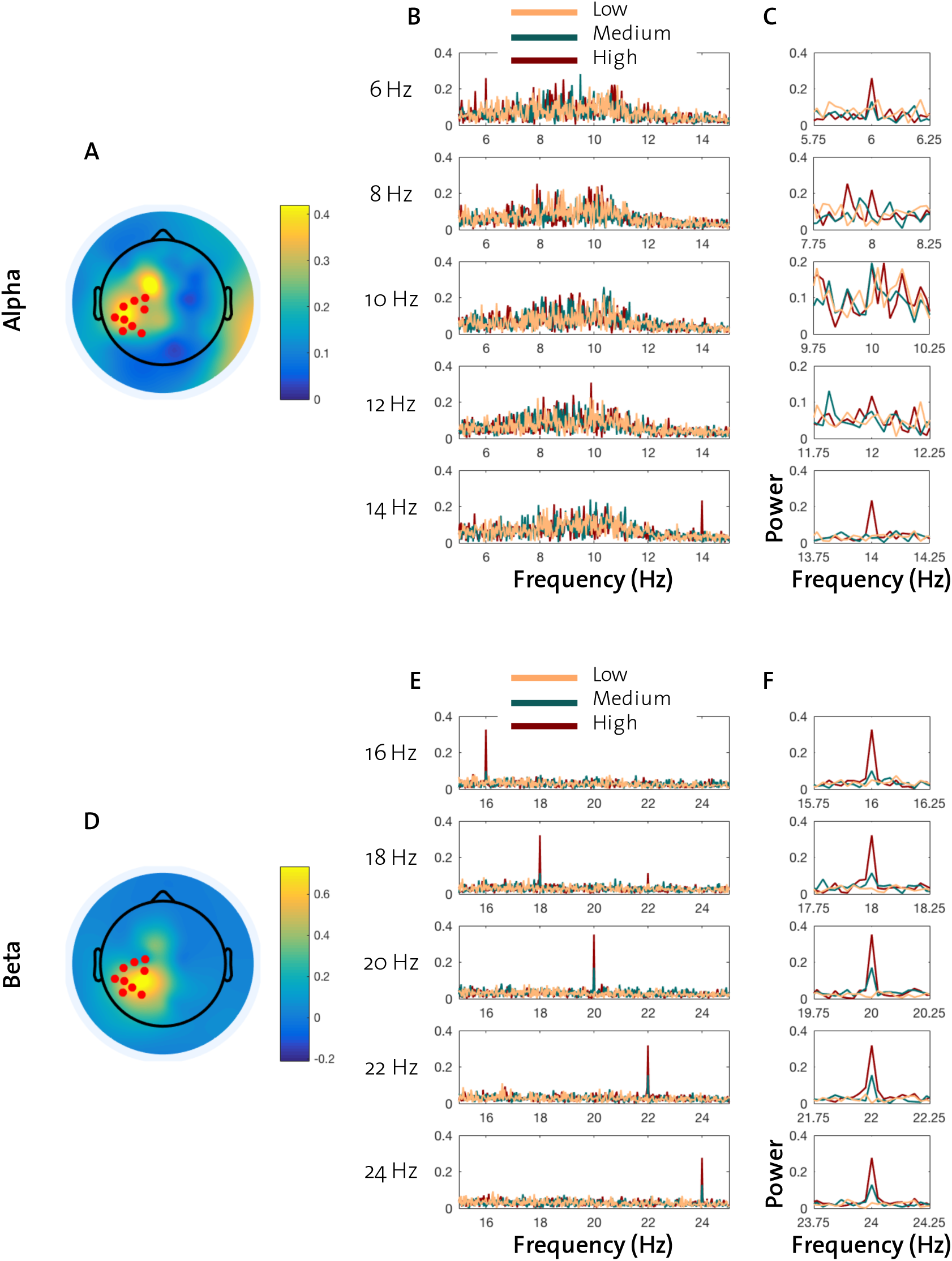
Power-based analysis of each stimulation condition averaged across subjects. A: Topographical plot of changes to electrode power across high-intensity alpha stimulations (red dots: ROI electrodes in the somatosensory area). B: Power spectra (FFT) of time-series of the selected electrode for stimulation frequencies in the alpha range. C: Closer depiction of power spectra at stimulation frequencies. D-F: Equivalent to A-C, across beta stimulation frequencies.

**Figure 4.**
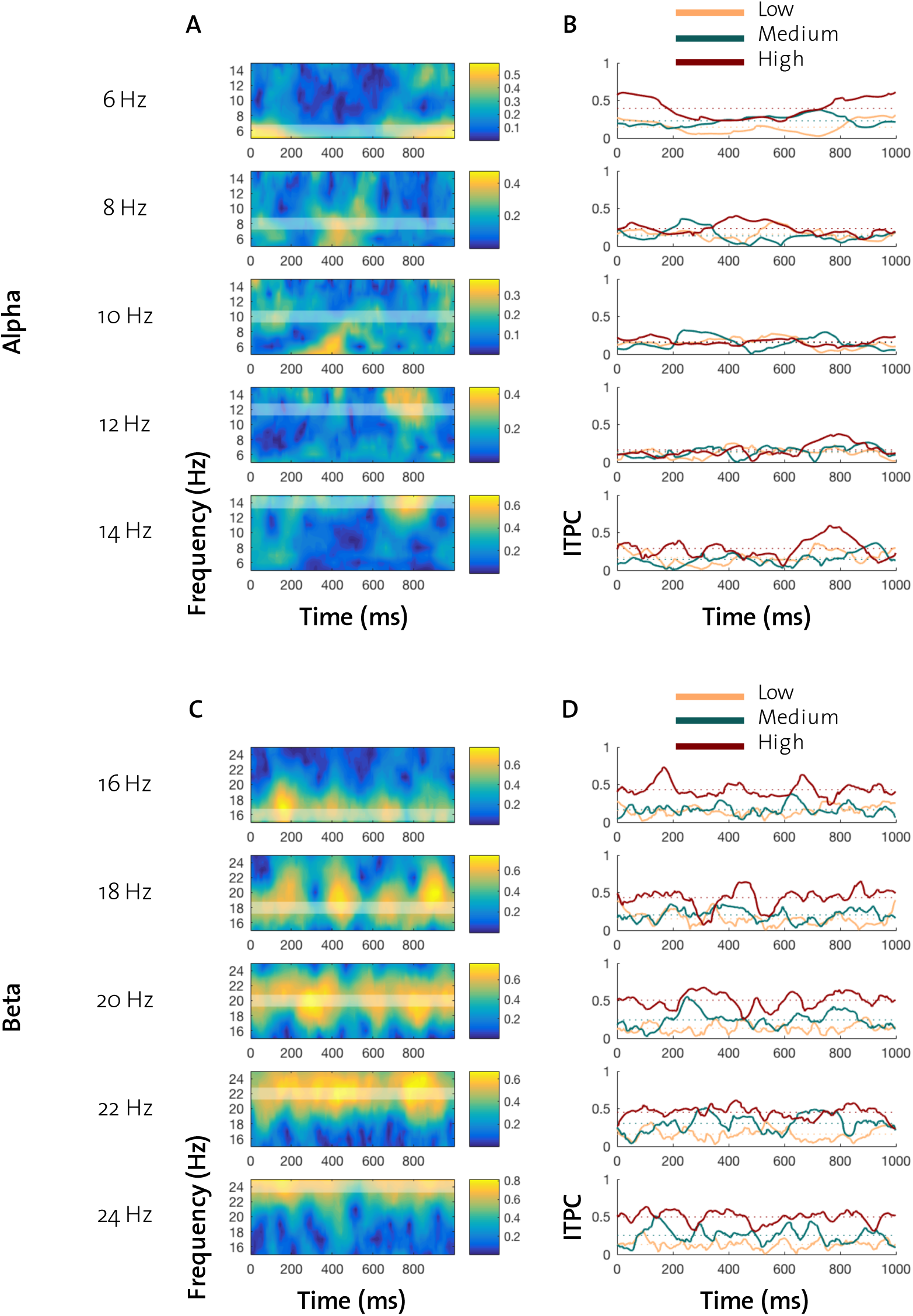
Phase-based analysis of each stimulation condition averaged across subjects. Time-series data from chosen electrodes (see Figure 3 A&D) were cropped in 1 s segments. A: Time-frequency plots of ITPC for stimulation frequencies delivered with high intensity in the alpha band with indication of the stimulation frequency (white transparent area). B: Time course of ITPC for each stimulation across alpha conditions. Dotted lines represent mean ITPC for each condition over the time course of the 1 second segment (averaged across trials). C&D: Equivalent to A&B, across beta stimulation frequencies.

**Figure 5.**
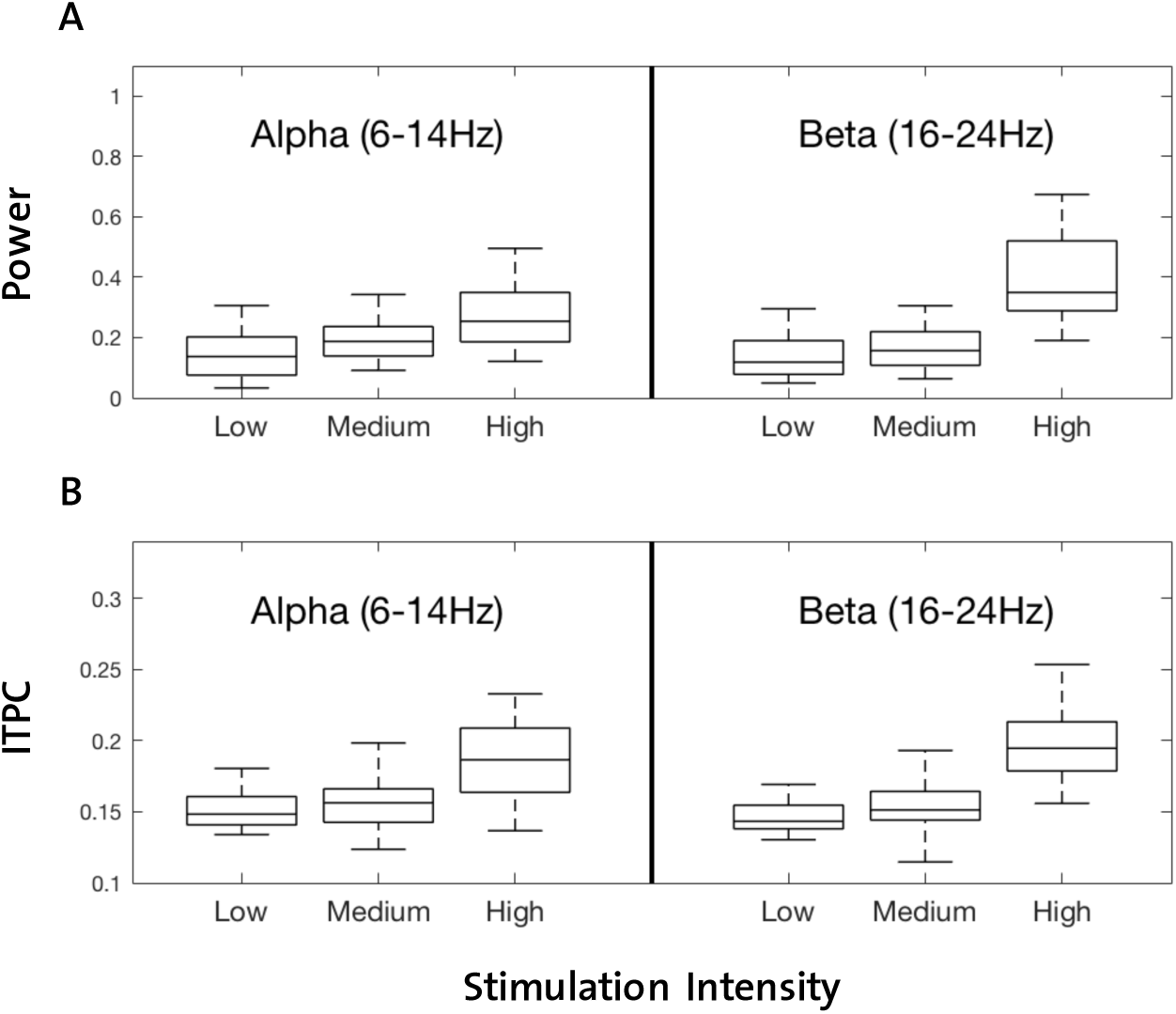
Differences of vibrotactile stimulation on frequency-specific power and ITPC across both frequency bands. A: Higher stimulation intensities reveal higher power values across the frequency bands. This main effect of intensity seems to be similar for alpha and beta (no interaction effect of frequency band and intensity). B: Also for ITPC we found a main effect of stimulation intensity. However, a significant interaction effect between frequency band and intensity suggests that the intensity effect differs between alpha and beta.

### ISPC in alpha and beta depend on stimulation intensity

To investigate entrainment effects of the stimulation signals, we computed changes to intersite phase clustering (ΔISPC) for each condition and used linear mixed effects models for the statistical analysis. Our model for the alpha band conditions included stimulation intensity and absolute distance from stimulation frequency to IAF (ΔIAF) as fixed factors. Any variations in slope and intercept of the data regarding individual variability was accounted for with the inclusion of subject as random variable. To test for an effect of stimulation intensity across all stimulation frequencies, we used mean-shifted ΔIAF values. Using the mean-shifted values allowed us to determine stimulation frequency-independent differences across the three intensity levels by directly comparing the resulting intercepts of the factor stimulation intensity. In line with our results from the power and ITPC analysis, we found the ΔISPC between stimulation and EEG signal to be more pronounced with increasing stimulation intensity. A post-hoc Tukey test revealed higher ISPC for the high intensity compared to sub-threshold and low (sub-threshold vs. high: p < 0.001; low vs. high: p = 0.004). However, the increase in ΔISPC from sub-threshold to low intensity showed no difference (p = 0.936). Testing the effect of stimulation intensity in the beta band, revealed similar results as seen for alpha. Again, linear mixed effects models were used with ΔISPC as dependent variable, stimulation intensity and mean-shifted distance from stimulation frequency to IBF (ΔIBF) as fixed variables and subject as random variable. Positive and statistically significant estimates for the post-hoc comparison between all intensity levels (sub-threshold vs. low: p < 0.001; sub-threshold vs. high: p < 0.001; low vs. high: p < 0.001), confirm the effect of stimulation intensity on steady-state neuronal response as hypothesized.

### Phase-coupling in beta depends on the interaction of intensity and distance to IBF

In order to test the main requirement regarding entrainment effects, and therefore the most important feature of the Arnold tongue, the dependency of stimulation effects on endogenous neuronal activity was investigated. We started with the analysis of the data obtained from stimulation conditions targeting alpha oscillations (stimulation frequencies from 6 to 14 Hz). The shape of the Arnold tongue suggests a dependency of entrainment effects (here: ΔISPC) on the distance between stimulation frequency to endogenous peak frequency (ΔIAF/IBF). In our statistical model this dependency can be depicted by plotting ΔISPC values according to their ΔIAF/IBF (see Figure 6 A/B). Negative slopes (derived for each stimulation intensity) which significantly differ from 0 can be expected, if the hypothesized dependency is true (i.e. stimulation frequencies closer to IAF/IBF lead to higher ΔISPC values). Although the slopes in the alpha conditions for low and high intensities show a negative direction (see Figure 6 A), statistically, they were not significantly different from 0 (ΔIAF:low: p = 0.998; ΔIAF:high: p = 0.915). Our next analysis focused on the beta band conditions (stimulation frequencies from 16 to 24 Hz). For beta, the slope of the low intensity differed significantly from 0 (p = 0.013), while sub-threshold and high intensities showed no such effect (ΔIBF:sub-threshold: p = 0.998; ΔIBF:high: p = 0.999). Our results from the beta band reveal a relationship between endogenous oscillations, stimulation intensity and phase coupling as would be assumed by the Arnold tongue. In our low intensity conditions, the distance of the stimulation frequency to IBF plays a crucial role in predicting coupling between stimulation and the underlying oscillation. The Arnold tongue widens by increasing the stimulation intensity, inferring a decreased effect of ΔIBF. This is because the system experiences such a strong enforcement that the ongoing oscillatory activity has no effect on ISPC.

**Figure 6.**
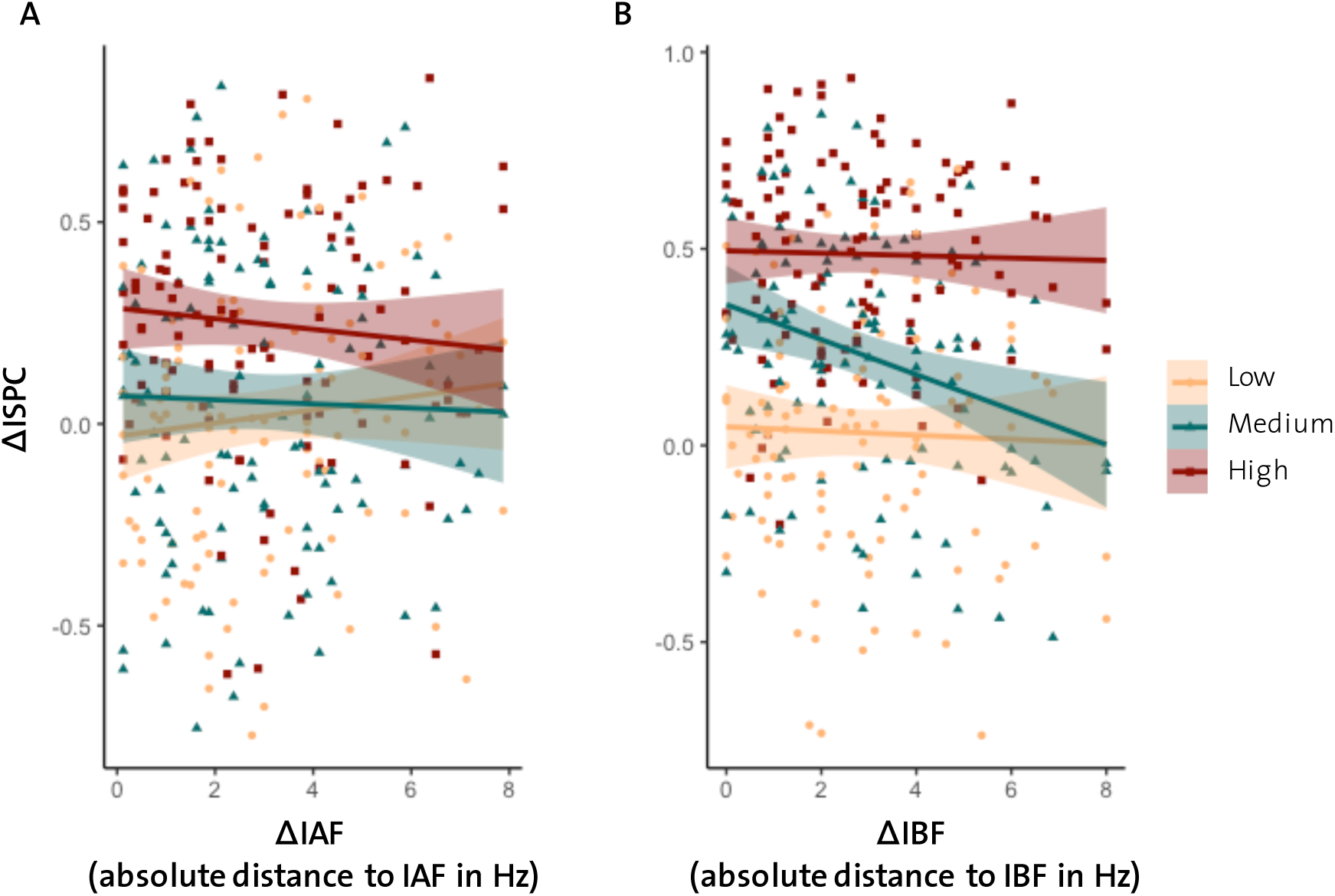
Depiction of the effects of different stimulation intensities on changes in ISPC in relation to the distance to endogenous frequencies (IAF and IBF). A: Stimulation conditions in the alpha band reveal no difference from 0 for any of the slopes and therefore no evidence for entrainment. B: In the beta band, low stimulation intensity reveals a negative slope which differs significantly from 0. This dependency of entrainment effect (ΔISPC) is in line with the hypothesized shape of the Arnold tongue.

## Discussion

In the present study, we investigated the effects of rhythmic vibrotactile stimulation across alpha and beta frequencies on neuronal activity in the somatosensory cortex. Our main results reveal a clear effect of stimulation intensity on frequency-specific power and intertrial phase clustering (ITPC) across both frequency bands. A significant interaction effect revealed a difference between alpha and beta stimulation frequencies with regard to their effect on frequency-specific power. The steeper power increase across the three stimulation intensities in beta compared to alpha, as well as the higher power values attained from high-intensity stimulation (Figure 5 A) is in line with previous findings suggesting that the resonance frequency of the somatosensory system is in the beta band (Müller et al., 2001; Snyder, 1992; Tobimatsu et al., 1999). Further, we provide evidence for the hypothesized entrainment effect by revealing a dependency of phase-locked activity between stimulation signal and endogenous beta brain oscillations on the interaction of stimulation intensity and frequency regarding its distance to individual beta peaks. This dependency is in line with the hypothesized Arnold tongue which predicts that entrainment via external signals of low intensity is stronger when the stimulation frequency matches the frequency of the endogenous system (Fröhlich, 2015; Notbohm et al., 2016; Pikovsky et al., 1999). Stimulation conditions in the alpha band revealed no such entrainment effect although effects of stimulation intensity on ISPC were found as well.

### Rhythmic sensory stimulation reveals online entrainment effects

Entrainment effects of sensory and non-invasive transcranial stimulation have been reported in various studies (for reviews see: Thut et al., 2011; Zoefel et al., 2018), yet our study is the first to demonstrate this effect for a wider range of endogenous brain oscillations in the somatosensory cortex. Our main finding that external rhythmic stimulation interacts with ongoing endogenous brain rhythms of the sensorimotor cortex is in line with previous reports using either sensory stimulation (Gulbinaite, van Viegen, Wieling, Cohen, & VanRullen, 2017; Notbohm et al., 2016) or various forms of rhythmic, non-invasive brain stimulation methods (Helfrich et al., 2014; Minami & Amano, 2017; Zaehle et al., 2010). However, until now entrainment effects have been mainly studied by targeting the visual alpha rhythm and its behavioral correlates. Previous studies revealed that steady-state evoked potentials depend on endogenous brain oscillations (Gulbinaite et al., 2017; Notbohm et al., 2016) and that modulating these endogenous rhythms of the visual cortex affects visual processing as measured at the behavioral level (Cecere, Rees, & Romei, 2015; Ronconi, Busch, & Melcher, 2018; Ronconi & Melcher, 2017). Together, these findings provide evidence for alpha entrainment effects in the visual system and also demonstrate the underlying causal relationship between brain oscillations and human behavior.

Few studies have investigated the modulation of neuronal oscillations in human somatosensory cortex. Two recent studies used transcranial alternating current stimulation (tACS) of the mu-rhythm (i.e. the alpha rhythm of sensorimotor cortex) to modulate sensorimotor perception threshold (Gundlach, Müller, Nierhaus, Villringer, & Sehm, 2016) and to reveal entrainment effects specific to endogenous mu-frequencies (Gundlach, Muller, Nierhaus, Villringer, & Sehm, 2017). In a first study, Gundlach et al. (2016) applied tACS stimulation at individual mu-alpha frequencies and revealed phase-dependent modulations of human somatosensory perception. However, in a tACS-EEG follow-up study, investigating the electrophysiological correlates of their tACS stimulation on endogenous mu-alpha oscillations, their findings reveal a counter-intuitive effect of a decrease of mu-alpha amplitude after stimulation (Gundlach et al., 2017). While the authors argue that homeostatic neuroplastic processes, as well as the electrode placement (bilateral) might have led to the decrease in mu-alpha, the underlying neural mechanism of their first study remains unclear. Even though their approach is well-suited to reveal functional modulations with tACS, it does not allow to investigate online entrainment effects due to large electrical artifacts. Further, due to the lack of different stimulation intensities, as well as variations in stimulation frequencies away from endogenous frequencies, the Arnold tongue cannot be derived from these studies. In addition, only mu-alpha frequency stimulation was applied, preventing conclusions in regard of other frequency bands. Here, we used tactile stimulation across various frequencies covering alpha and beta frequencies, which was provided mechanically and, therefore, did not interfere with the EEG recordings, thus, enabling the analysis of potential entrainment effects during stimulation. Our results suggest that rhythmic activity in somatosensory cortex can be influenced, but that potential entrainment effects were only found in the beta band.

### Stronger response to rhythmic beta stimulation in the somatosensory system

Although studies investigating spontaneous (resting-state) EEG activity, found both beta (mu-beta) and alpha (mu-alpha) oscillations in sensorimotor areas (Hillebrand, Barnes, Bosboom, Berendse, & Stam, 2012; A. Keitel & Gross, 2016; Ramkumar, Parkkonen, & Hyvarinen, 2014), the steady-state evoked potentials in the two frequency bands differ clearly (see Figure 3). This finding is in line with previous studies using rhythmic sensory stimulation, revealing that each sensory system responds maximally to a specific stimulation frequency (for a review: Vialatte et al., 2010). Arguably, this represents a mechanism by which each of the sensory systems processes information optimally (Hutcheon & Yarom, 2000). Such a temporal tuning function appears in the visual system around 10 Hz (Regan, 1989), in the auditory system around 40 Hz (Stapells, Makeig, & Galambos, 1987) and in the somatosensory system around 21 Hz (Müller et al., 2001; Snyder, 1992; Tobimatsu et al., 1999; Tobimatsu, Zhang, Suga, & Kato, 2000). Such resonance mechanisms could be responsible for our mixed results for alpha and beta stimulation frequencies.

### True entrainment or rhythmically evoked potentials?

Our results are in line with previously stated dependencies of EEG responses to rhythmic stimulation on endogenous brain oscillations. However, it is difficult to draw a final conclusion on whether steady-state somatosensory evoked potentials (SSSEPs), as produced in our study with vibrotactile stimulation, represent entrained neural oscillations. As Haegens and Zion-Golumbic (2018) point out in a recent review, the interpretation of neuronal responses to rhythmic stimulation as neural entrainment might lack the fundamental distinction from a rhythmic series of evoked potentials. Although some studies have shown that non-rhythmic (= jittered) stimulations do not reveal a non-linear response with dependency on endogenous oscillations (Kayser, Ince, Gross, & Kayser, 2015; Notbohm et al., 2016), others found the opposite effect (Capilla et al., 2011; C. Keitel, Thut, & Gross, 2017; see also: Zoefel et al., 2018). Our findings can be seen in agreement with an entrainment mechanism, as we find dependencies of SSSEPs on endogenous oscillations only with a low stimulation intensity and only in beta. Arguably, if SSSEPs would in fact represent only regular repetitions of evoked potentials, we should have seen the same effects for alpha and beta stimulation. However, we do not evaluate our finding in comparison to non-rhythmic stimulation. Regarding the ongoing debate on whether neural entrainment occurs in response to rhythmic sensory stimulation, modulating functional correlates of neural oscillations (e.g. perception, memory) would serve as a convincing outcome measure (Haegens & Zion Golumbic, 2018). Further, in the present study, we used sensory data from selected electrodes that revealed strongest frequency-specific power changes in a finger tapping task prior to the main experiment. Although our data reveal clear peaks in the alpha and beta bands for most of the participants, our analyses have to be interpreted with caution. For example, a decline in attention and alertness throughout the experiment could have led to variations in IAF and/or IBF (Mierau, Klimesch, & Lefebvre, 2017), reducing potential entrainment effects as analyzed here. Nevertheless, our findings provide valuable insight in the neural processing of rhythmic sensory stimulation and should be considered in future studies using somatosensory entrainment.

## Conclusion

The concept of modulating brain oscillations by rhythmic stimulation has become increasingly popular in research on brain rhythms and their underlying functions (Thut et al., 2011). Despite the limitations of the present study, we show that there is a significant frequency-specific dependency of brain responses to rhythmic vibrotactile stimulation on individual neuronal activity in the beta band. We conclude that our findings support the notion of entrainment of endogenous brain oscillations in somatosensory cortex. This opens up possibilities to causally test interactions between oscillatory activity and behavior by utilizing entrainment via somatosensory stimulation.

## Acknowledgments

We thank Pearl Saldanha and Marionna Münger for their help with data acquisition, and the Statistical Counseling Service of ETH Zurich, especially Janine Burren, for methodological support. We also acknowledge the support of the Neuroscience Center Zurich (ZNZ).

## Notes

**Conflict of Interest** The authors declare that the research was conducted in the absence of any commercial or financial relationships that could be construed as a potential conflict of interest.

**Funding** This research was funded by the Swiss National Science Foundation (No. 320030_175616).

